## Supplementary Figures and Methods for "Prediction of designer-recombinases for DNA editing with generative deep learning"

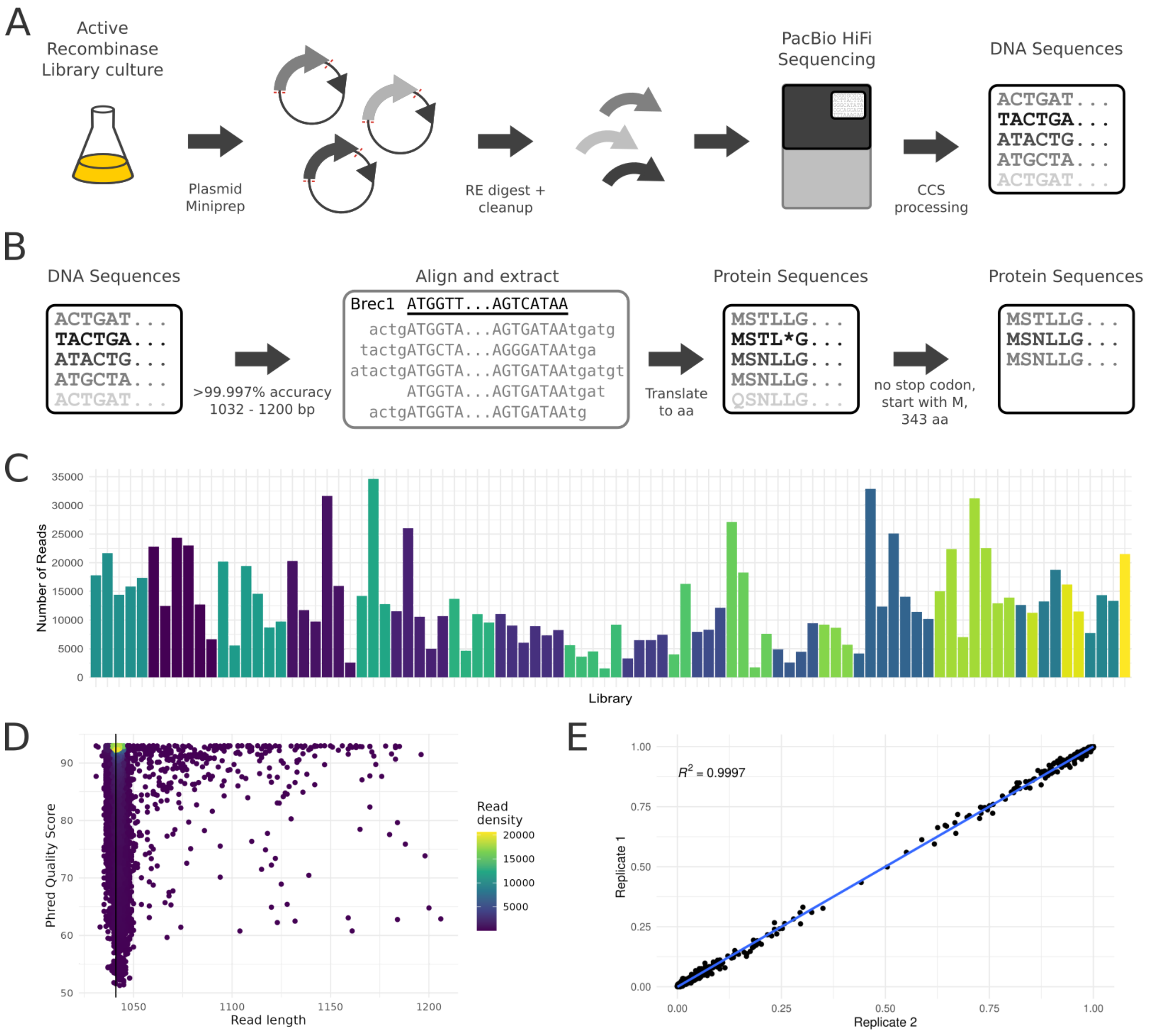

**Supplementary Figure 1.** Sequencing pipeline scheme and sequence data properties. **A)** Recombinase library sequencing preparations. Plasmids encoding active recombinases are extracted from *E. coli* culture. Recombinase genes are then cut by restriction enzyme digestion and enriched by SPRI-bead cleanup. Sequencing of the genes is then performed on a PacBio Sequel with the HiFi sequencing approach. The acquired subreads were then processed with ccs to generate high quality consensus sequences. **B)** Processing of sequencing data. DNA sequences are filtered for accuracy and length. Gene sequences are then isolated by alignment to Brec1 (Karpinsky et al. 2016) and converted to amino acid. Finally, resulting protein sequences are filtered for correct length, sequences that start with methionine, and sequences that do not contain a stop codon in the middle of the sequence. **C)** Total number of recombinase sequences acquired per Library, colored by evolution project. **D)** Example analysis of loxE-3 DNA read length and read quality acquired by PacBio HiFi sequencing. Calculated sequencing quality is beyond the required 99.997 % accuracy (Phred Score of ~25). **E)** Correlation of repeated sequencing of loxE-1. Replicate 1 and Replicate 2 were prepared from independent plasmid extractions from separate cultures and sequenced on different sequencing runs. Each point represents the observed frequencies of the amino acids on each residue position of the proteins. Replicate sequences are highly correlated, $R^2$ was calculated as 0.9997.

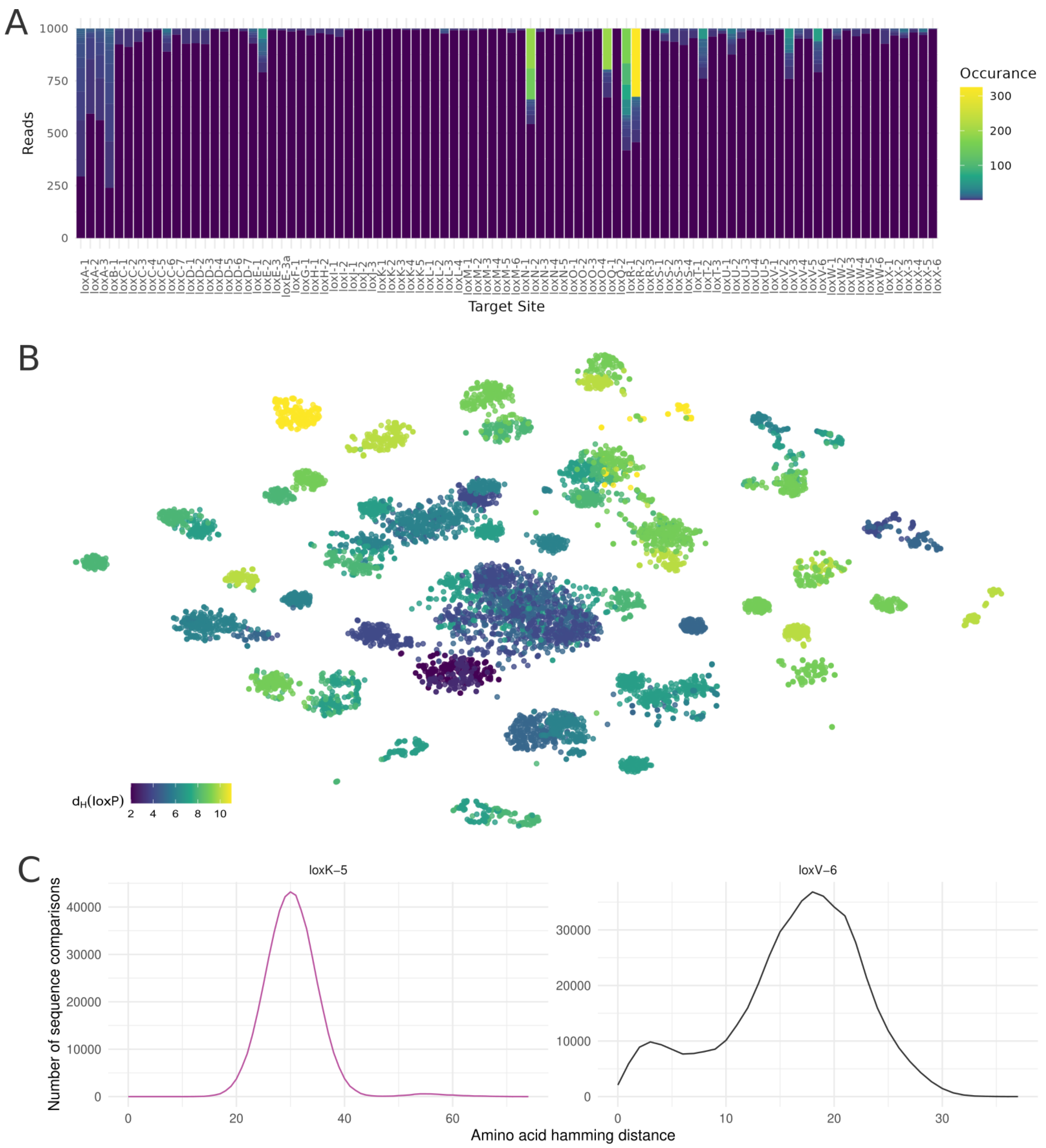


**Supplementary Figure 2.** Sequence data composition. **A)** Number of sequence reads occurring multiple times per library. The color code depicts sequence occurrence. Note the high diversity of clones within most libraries. **B)** t-SNE dimensionality reduction of 100 recombinase sequences from all sequenced libraries. Color indicates the base pair hamming distance (dH(loxP)) of the associated target sites to the loxP sequence. **C)** All vs. all amino acid sequence hamming distances of loxK-5 and loxV-6 libraries. Comparisons were made with 1000 sequences per library.

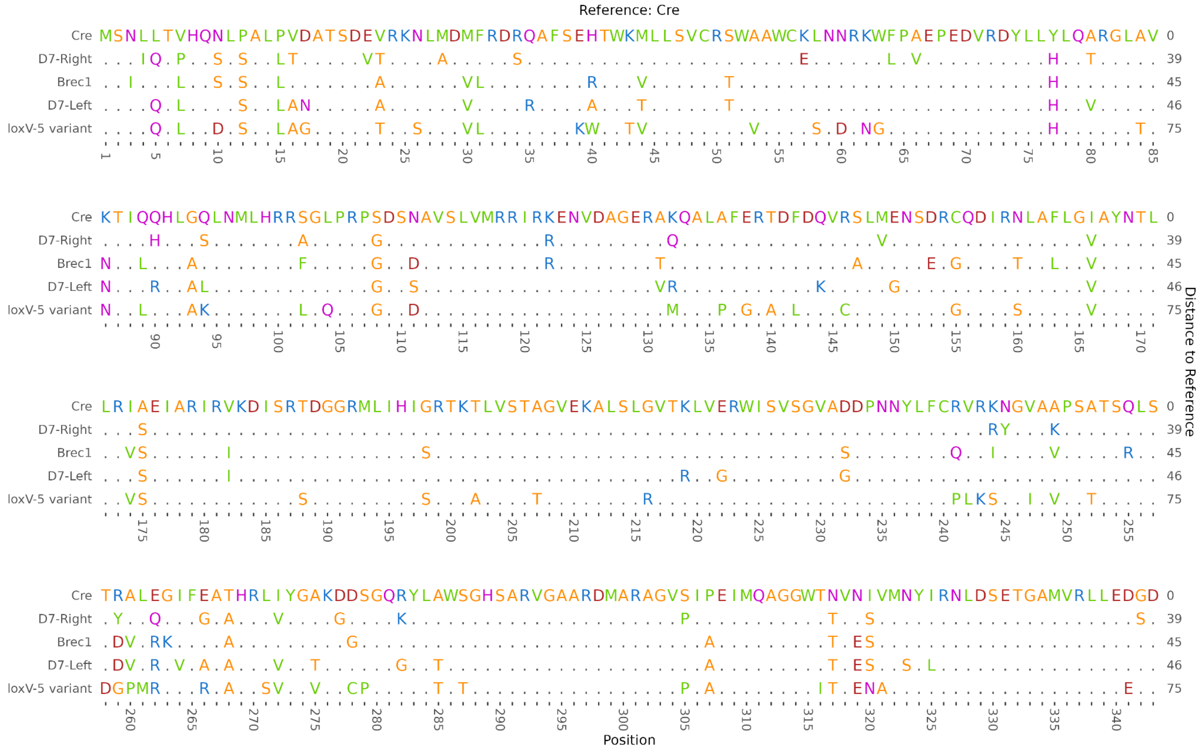

**Supplementary Figure 3.** Alignment of recombinase sequence with highest observed difference to Cre. D7-Left, D7-Right (Lansing et al. 2022) and Brec1 (Karpinsky et al. 2016) are included for comparison.

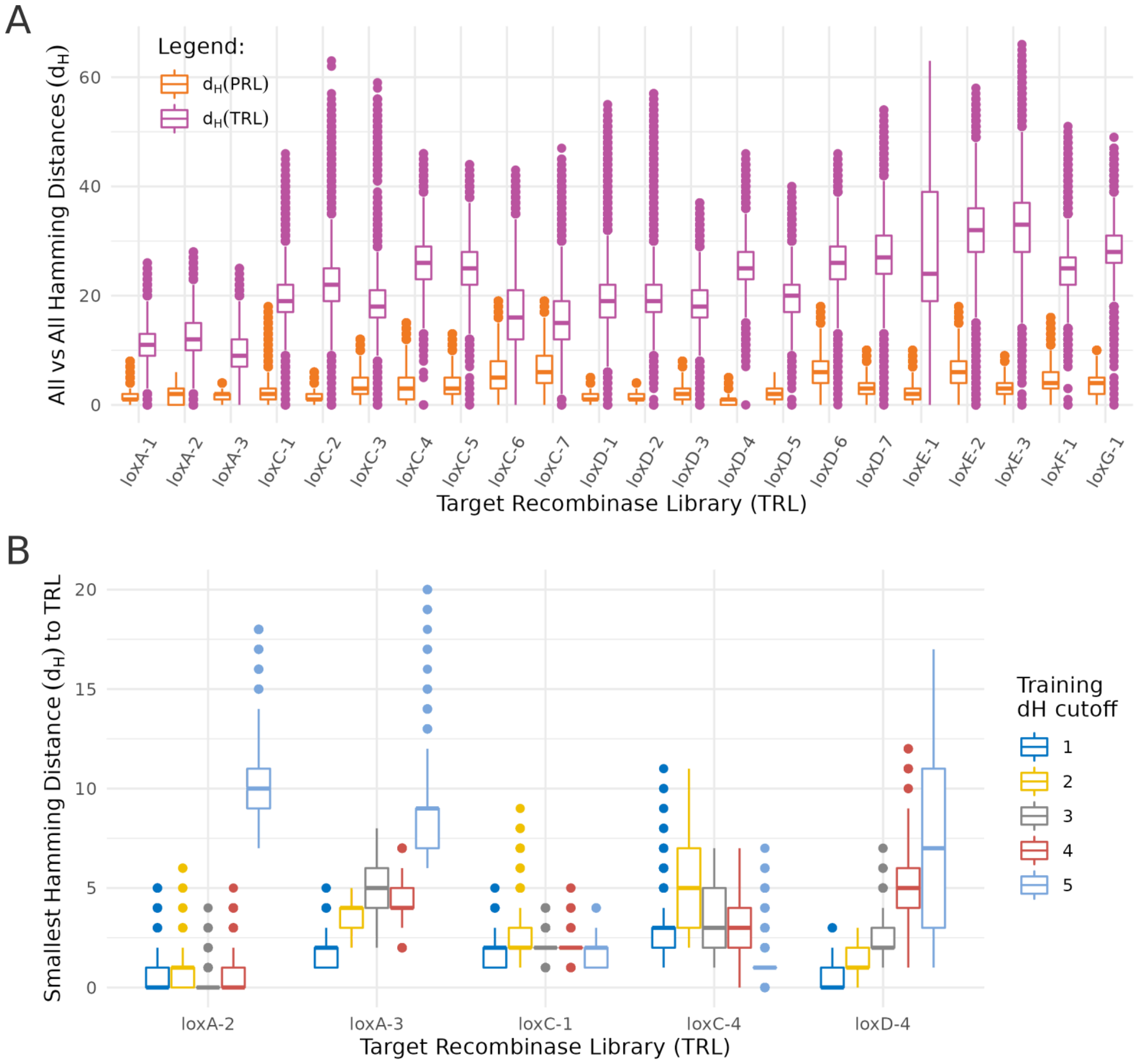


**Supplementary Figure 4. A)** Sequence variance of selected TRL and LOOCV PRL. Variance of the sequences was determined by all vs. all sequences comparison. Distance is measured in hamming of the amino acid sequences. TRL has a much higher variance than PRL in all libraries. **B)** LOOCV prediction of selected TRL with increasing number of neighboring libraries removed from training data. Training dH cutoff defines the minimum base pair distance to the TRL target site still allowed in the training data. Increasing the cutoff value causes the PRL to be less similar to the TRL in some samples, while others are less affected.

### Depletion of plasmid backbone with SPRI-beads

We followed dx.doi.org/10.17504/protocols.io.n7hdhj6 to make custom SPRI-beads with Tween. However, instead of using 0.4% Sera-Mag SpeedBeads Carboxyl Magnetic Beads (GE Healthcare) we used 2%. Additionally, we also made a concentrated SPRI-beads mix. We took 50ul Sera-Mag beads and washed them 3 times with nuclease free water, followed by complete removal of the liquid. To this we added a version of the custom SPRI-bead mix without the Sera-Mag beads and nuclease free water so that a mixture of beadless custom SPRI-bead mix and water with Sera-Mag beads equals 0.75x to 1 in a total volume of 50 µl.

To deplete the pEVO plasmid backbone, a fragment of ~5 kb size, we mix 0.75x custom SPRI-bead mix with 1x of the XbaI + BsrgI digested pEVO. After 15 minutes of incubation we placed the mixture on a magnet and transferred the supernatant to a new microcentrifuge tube. To this new tube we then mixed in 2 µl of the concentrated beads and incubated for 3 minutes. The supernatant is then transferred again and another 2 µl of concentrated beads is added. The liquid at this stage almost exclusively contained the recombinase fragments. We transferred the liquid to another tube and mixed in 0.35x Ampure XP (Beckman Coulter) relative to the volume of our supernatant. After incubating for 15 min we performed a normal Ampure XP bead cleanup with 2x 75% EtOH wash and an elution with 15 µl nuclease free water.
